# Feeding Experimentation Device version 3 (FED3): An open-source home-cage compatible device for measuring food intake and operant behavior

**DOI:** 10.1101/2020.12.07.408864

**Authors:** Bridget A. Matikainen-Ankney, Thomas Earnest, Mohamed Ali, Eric Casey, Amy K. Sutton, Alex Legaria, Kia Barclay, Laura B. Murdaugh, Makenzie R. Norris, Yu-Hsuan Chang, Katrina P. Nguyen, Eric Lin, Alex Reichenbach, Rachel E. Clarke, Romana Stark, Sineadh M. Conway, Filipe Carvalho, Ream Al-Hasani, Jordan G. McCall, Meaghan C. Creed, Victor Cazares, Matthew W. Buczynski, Michael J. Krashes, Zane Andrews, Alexxai V. Kravitz

## Abstract

Feeding is critical for survival and disruption in the mechanisms that govern food intake underlie disorders such as obesity and anorexia nervosa. It is important to understand both food intake and food motivation to reveal mechanisms underlying feeding disorders. Operant behavioral testing can be used to measure the motivational component to feeding, but most food intake monitoring systems do not measure operant behavior. Here, we present a new solution for monitoring both food intake and motivation: The Feeding Experimentation Device version 3 (FED3). FED3 measures food intake and operant behavior in rodent home-cages, enabling longitudinal studies of feeding behavior with minimal experimenter intervention. It has a programmable output for synchronizing behavior with optogenetic stimulation or neural recordings. Finally, FED3 design files are open-source and freely available, allowing researchers to modify FED3 to suit their needs. In this paper we demonstrate the utility of FED3 in a range of experimental paradigms.

**In Brief:** Using a novel, high-throughput home cage feeding platform, FED3, Matikainen-Ankney et al. quantify food intake and operant learning in groups of mice conducted at multiple institutions across the globe. Results include rates of operant efficiency, circadian feeding patterns, and operant optogenetic self-stimulation.

**Highlights:**

- The Feeding Experimentation Device version 3(FED3) records food intake and operant behavior in rodent home cages.
- Analysis of food intake includes total intake, meal pattern analysis, and circadian analysis of feeding patterns.
- FED3 also allows for operant behavioral assays to examine food learning and motivation.

## Introduction

Feeding is critical for survival and dysregulation of food intake underlies medical conditions such as obesity and anorexia nervosa. Quantifying food intake is necessary for understanding these disorders in animal models. However, it is challenging to accurately measure food intake in rodents due to the small volume that they eat. Researchers have devised multiple methods for quantifying food intake in rodents, each with advantages and drawbacks (Ali and Kravitz, 2018). Traditional methods rely on weighing differences in food pellets across hours or days. This is time consuming to complete, can be subject to error and variability, and does not allow for fine temporal analysis of consumption patterns (Acosta-Rodríguez et al., 2017; Reinert et al., 2019). Automated tools have been developed for measuring food intake in home cages with high temporal resolution, although most require modified caging, powdered foods, or connected computers that limit throughput and are incompatible with vivarium environments (Ahloy-Dallaire et al., 2019; Farley et al., 2003; Moran, TH, 2003; Yan et al., 2011).

In addition to measuring total food intake, understanding neural circuits involved in feeding requires exploring *why* animals seek and consume food. Has their motivation for a specific nutrient changed? Has their feeding gained a compulsive nature that is insensitive to satiety signals? These questions can be answered with operant tasks, where rodents receive food contingent on pressing levers or activating “nose-pokes” (Curtis et al., 2019; Mourra et al., 2020; O’Connor et al., 2015; Wald et al., 2020). Typically, operant behavior is tested in dedicated chambers, where animals are tested for a few hours each day. Training in dedicated operant chambers has several limitations: tasks can take weeks for animals to learn, animals are often tested at different phases of their circadian cycle due to equipment availability, and food restriction can be necessary to get animals to seek food in the operant chamber, which is a confound in feeding studies. To mitigate these issues, researchers have begun to test operant behavior in rodent home cages, resulting in both fewer interventions from the researcher and faster rates of learning (Balzani et al., 2018; Francis and Kanold, 2017; Lee et al., 2020).

Here, we present a new solution for monitoring both food intake and operant behavior in rodent home cages: the Feeding Experimentation Device version 3 (FED3). Our goal was to develop a device for measuring food intake in rodent home cages with high temporal resolution, while also measuring food motivation via operant behavior. FED3 contains a pellet dispenser, two “nose-poke” sensors for operant behavior, visual and auditory stimuli, and a screen for experimenter feedback. FED3 is compact and battery powered, fitting in normal vivarium home-cages without any connected computers or external wiring. FED3 also has a programmable output that can control other equipment, for example to trigger optogenetic stimulation after a nose-poke or pellet removal, or to synchronize feeding behavior with electrophysiological or fiber photometry recordings. Finally, FED3 is open-source and was designed to be customized and re-programmed to perform novel tasks to help researchers understand food intake and food motivation. Here, we describe the design and construction of FED3, and present several experiments that demonstrate its versatile functionality. These include measuring circadian patterns of food intake over multiple days, performing meal pattern analysis, automated operant training, and optogenetic self-stimulation. FED3 extends existing methods for quantifying food intake in rodents and can help researchers achieve a deeper understanding of feeding and feeding disorders.

## Results

### Hardware and design of the Feeding Experimentation Device version 3 (FED3)

The Feeding Experimentation Device version 3 (FED3) is controlled by a 48mHz ATSAMD21G18 ARM Cortex M0 microprocessor (Adafruit Adalogger M0) and contains two nose-pokes, a pellet dispenser, a speaker for auditory stimuli, 8 multi-color LEDs for visual stimuli, a screen for experimenter feedback, and a programmable analog output (Figure 1A, B). When mice interact with FED3, the timing of each poke and pellet removal are logged to an internal microSD card for later analysis and summary data is displayed on the screen for immediate feedback to the researcher (Figure 1B). FED3 is small (~10cm x 12cm x 9cm), battery-powered, and completely self-contained so it fits in standard vivarium home-cages without modification or introducing wiring to the cage (Figure 1C). It is powered by a rechargeable battery that lasts ~1 week between charges (exact battery life can depend on the behavioral program). FED3 also has magnetic mounts to facilitate wall-mounting on any plastic box to mimic a traditional operant setup (see Supplementary Video 1). We provide programs for running rodents on multiple common behavioral paradigms, including free feeding, time-restricted free-feeding, fixed-ratio (FR1), and progressive ratio (PR) operant tasks, and have written a FED3 Arduino library to ease development of custom programs. FED3 also has a programmable output that allows synchronization with external equipment for aligning behavioral events with fiber-photometry recordings (London et al., 2018; Mazzone et al., 2020), electrophysiological recordings (London et al., 2018), or other equipment such as video tracking systems (Krynitsky et al., 2020; Li et al., 2019). Finally, we provide a graphical analysis package written in Python that enables users to generate detailed plots from FED3 data (Figure 2F). FED3 is open-source and freely available online, including 3D design files, printed circuit board (PCB) files (Figure 1D), build instructions (Figure 1E), and code (https://github.com/KravitzLabDevices/FED3).

**Figure 1.**
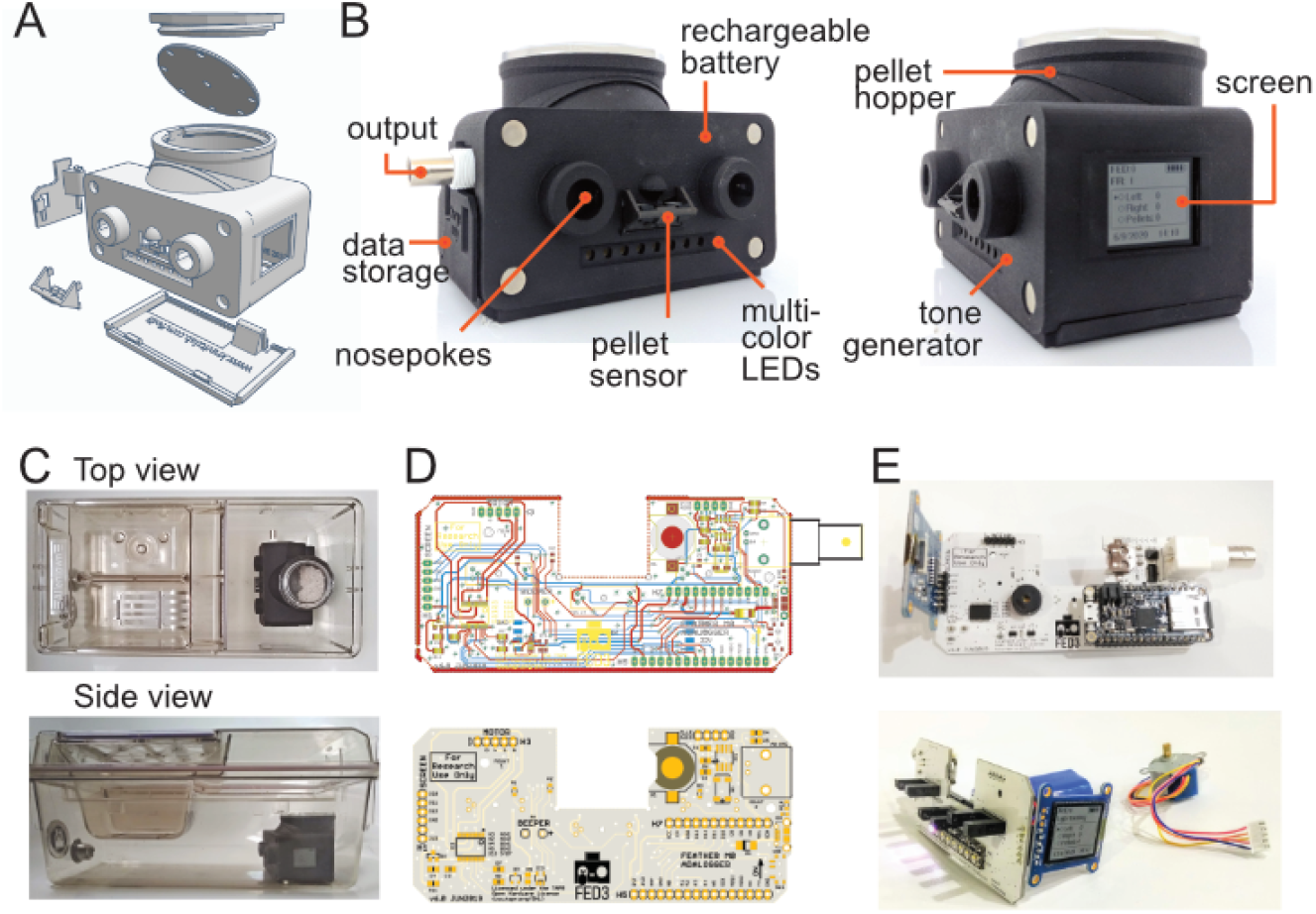
Assembly of FED3. **(A)** Exploded view schematic, and **(B)** photos of FED3 with main components highlighted. **(C)** Assembled FED in an Allentown NextGen home-cage, top view, and side view. **(D)** Schematic view of printed circuit board (PCB, top). Rendering of the PCB (bottom). **(E)** Back view (top) of assembled and populated FED3 PCB, and side view (bottom) of assembled FED3 electronics.

**Figure 2.**
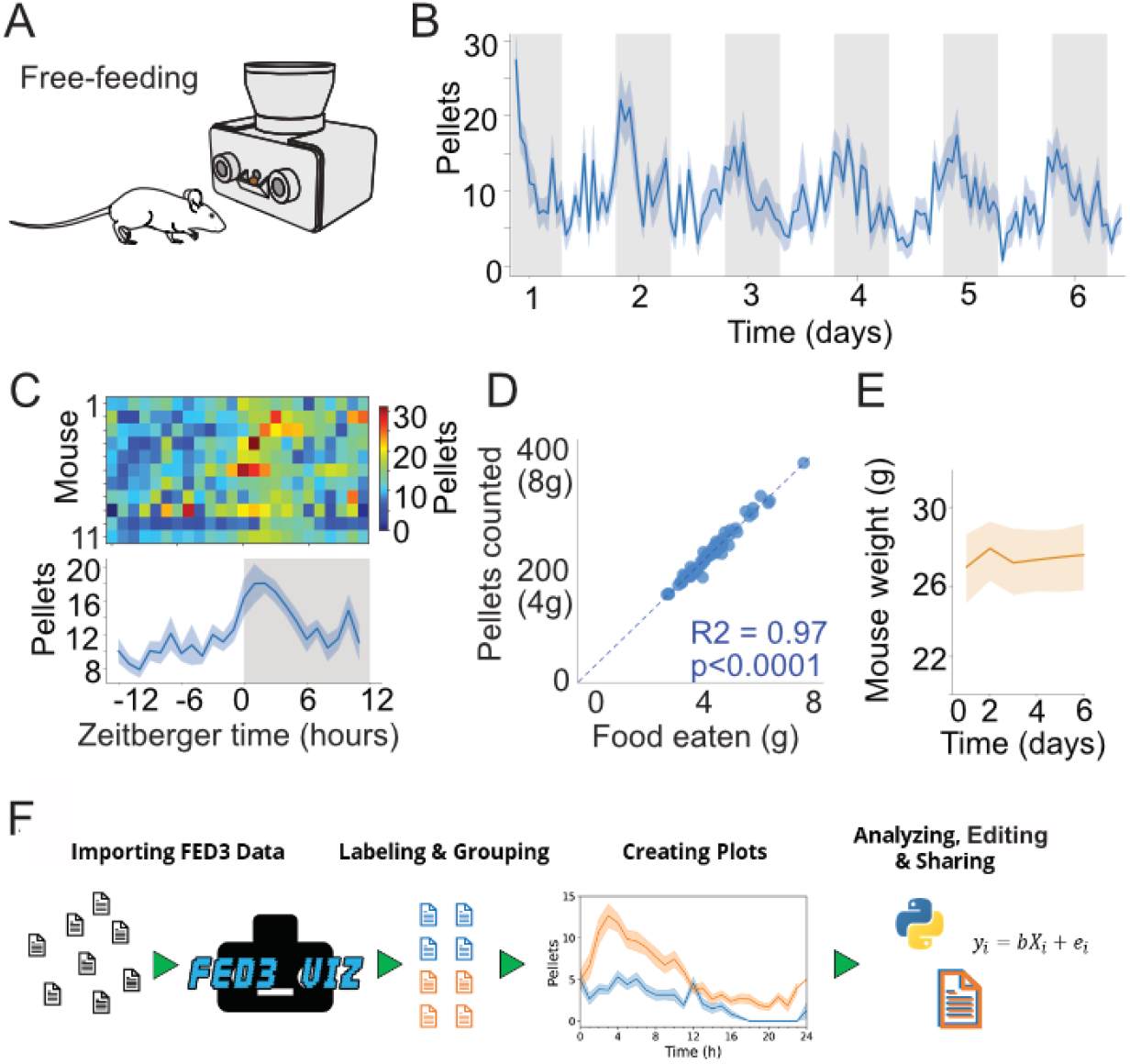
FED3 tracks food intake. **(A)** Schematic of FED3 in free-feeding mode. **(B)** Pellets per hour across 6 consecutive days. Shaded areas indicate dark cycle. **(C)** Chronograms of pellets eaten in heatmap (top) and line plot. **(D)** Regression of the calculated weight of the pellets recorded by FED vs the measured difference in weight of the FED between days. **(E)** Mice weight across time during free feeding. **(F)** Schematic of FED3VIZ workflow. Data is shown as means ± SEM in **(B, C, E)**.

### Quantifying total food intake

To demonstrate how FED3 can be used for quantifying food intake, eleven FED3 devices were set to the “free-feeding” program, which begins by dispensing a pellet and monitoring its presence in the pellet well (Figure 2A). Each time the pellet is removed, FED3 logs the date and time to the storage card and dispenses another pellet. This paradigm allows for the reconstruction of detailed feeding records over multiple days, and knowing the caloric content of the pellets enables understanding of caloric intake. Devices were placed with singly housed mice for 6 consecutive days with no additional food source, resulting in the expected circadian rhythm in food intake (Figure 2B, C). To test accuracy, we confirmed that the daily change in weight of the device correlated linearly with the number of pellets removed multiplied by the weight of each pellet (20mg, Figure 2D). The coefficient of determination (R^2^) for the regression was 0.97, indicating an error rate of ~3% between manual weighing and counting of pellets by FED. Finally, we confirmed that the mice obtained all of their necessary daily calories from FED3, evidenced by a stable body weight across these six days (Figure 2E). In other experiments, we have run FED3 for >1 month without observing weight loss.

FED3 creates a large amount of data, particularly when run over multiple days. To facilitate analysis of FED3 data, we created FED3VIZ, an open-source Python-based analysis program (https://github.com/earnestt1234/FED3_Viz). FED3VIZ offers plots for visualizing different aspects of FED3 data, including pellet retrieval, poke accuracy, pellet retrieval time, delay between consecutive pellet earnings (inter pellet intervals), meal size, and progressive ratio breakpoint. These metrics can be plotted for single files or group averages. FED3VIZ’s averaging methods provide options for aggregating data recorded on different days, while preserving time-of day or phase of the light-dark cycle. Furthermore, circadian activity patterns can also be visualized with the FED3VIZ “Chronogram” and “Day/Night Plot” functions, which segment data based on a user-defined light cycle. The processed data going into each plot can also be saved and used to create new visualizations or compute statistics in other programs, promoting reproducible, sharable analysis and visualization of FED3 data.

### Meal analysis

FED3 records the date and time that each pellet is removed, which can be used to quantify feeding patterns including the size and quantity of meals, eating rate within meals, and timing between meals. Different parameters have been used by different groups to define meals, often based on numerical cut-offs (Farley et al., 2003; Kanoski et al., 2013; Melhorn et al., 2010). To complement these approaches, we aimed to develop an unbiased approach for understanding meal patterns, based on the distribution of time intervals between each pellet consumed. For the eleven mice in the above experiment, we plotted the inter-pellet time interval histogram (Figure 3A) for both the dark and light cycle, based on an approach that was previously established in rats (Cottone et al., 2007). We observed a large peak at <1min between pellets, meaning that the vast majority of pellets in this study were consumed within 1-min of another pellet. We used this distribution to classify pellets eaten within the same minute as belonging to the same meal, and pellets with larger intervals between them belonging to different meals. This approach revealed that animals eat fewer meals during the light cycle (Figure 3B), but eat similar number of pellets within each meal and obtain a similar fraction of their pellets within meals in each cycle. Differences in meal patterning have been linked to obesity (Farley et al., 2003; Wald and Grill, 2019), so this analytical approach may assist in understanding obesity and other disorders of feeding.

**Figure 3.**
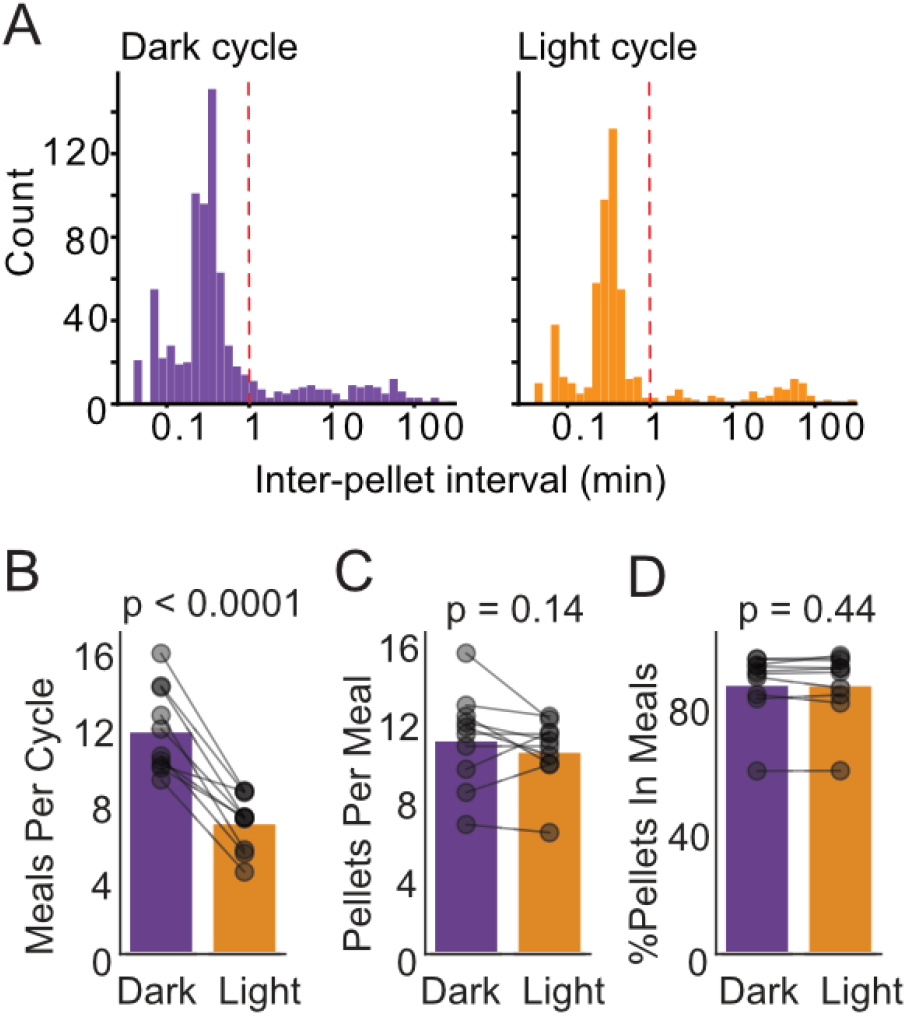
Meal analysis with FED3. **(A)** Inter-pellet interval histograms for Dark and Light cycle feeding. **(B)** Meals per day **(C)** Pellets per meal and **(D)**% of pellets within meals for Dark and Light cycle feeding. N=10 mice, paired t-tests.

### Circadian patterns of operant behavior

A unique feature of FED3 is that it is small and wire-free, allowing it to fit inside of traditional vivarium home-cages. This facilitates quantification of circadian rhythms in both operant and feeding behavior (Figure 4A). To demonstrate this capability, we quantified how nose-poking varied over the circadian cycle, by running mice on a fixed-ratio 1 (FR1) task for six consecutive days. We tracked active (correct) and inactive (incorrect) pokes. As expected, both active and inactive pokes were higher during the dark cycle (Figure 4B-D). Surprisingly, however, poke *accuracy* was slightly (higher during the light cycle (Figure 4E), suggesting that operant responding is more efficient during the light cycle.

**Figure 4.**
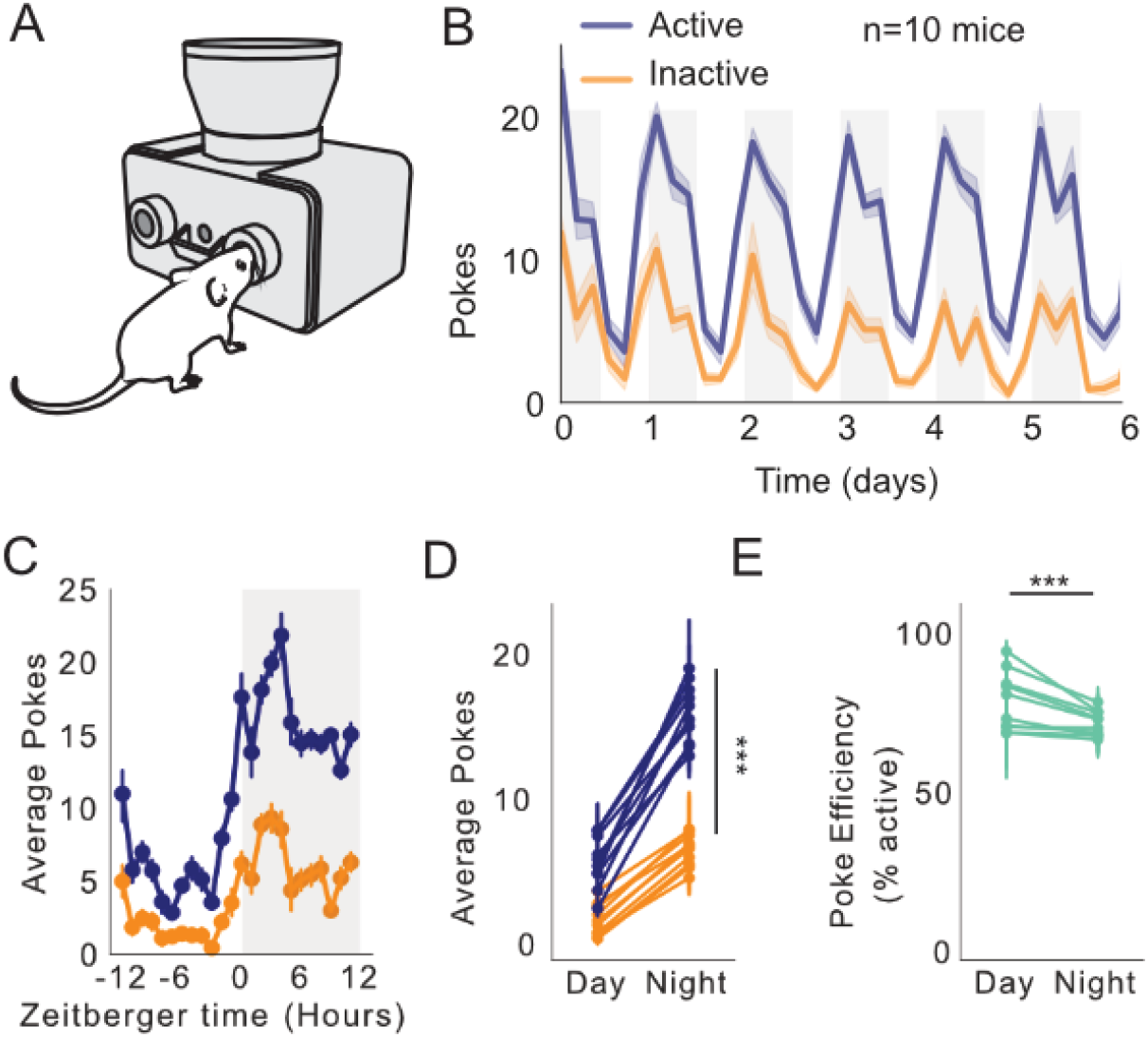
FED3 reveals circadian feeding patterns. **(A)** Schematic of FED3 in FR1 mode. **(B)** Active (blue) and inactive (orange) pokes over six days. n=10 mice. **(C)** Average pokes (active, blue; inactive, orange) over 24 hour cycle. **(D)** Average pokes during light and dark cycles. Significant interaction between day/night and average pokes (F(1,792)=85.225, p<0.0001); significant effect of day/night (F(1,792)=668.700, p<0.0001); significant effect of active or inactive pokes (F(1,792)=610.824, p<0.0001). Fitted linear model for two-way ANOVA, (F(3,792)=454.917, p<0.0001.) **(E)** Average poke efficiency (p=0.0001, student’s t-test).

### A multi-site study of learning rates with FED3

Multiple research groups currently use FED3. We therefore asked how mouse operant learning varies across different laboratories. To do this, we obtained operant data from the first overnight FR1 session from seven laboratories, resulting in data from 122 mice (mix of males and females, Figure 5A). When comparing pellets earned in an overnight session across laboratories, we observed a significant effect of group, linked to significant differences in 4 out of 21 post-hoc comparisons (Figure 5B). When viewed as a single distribution (Figure 5B inset), the distribution of pellets earned from all groups was consistent with a single Gaussian distribution (p>0.05, Shapiro-Wilk test for normality). This highlights how FED3 enables high throughput studies of operant behavior, and also demonstrates the potential for false positive effects when comparing between groups with small sample sizes (Button et al., 2013) (Figure 5B).

**Figure 5.**
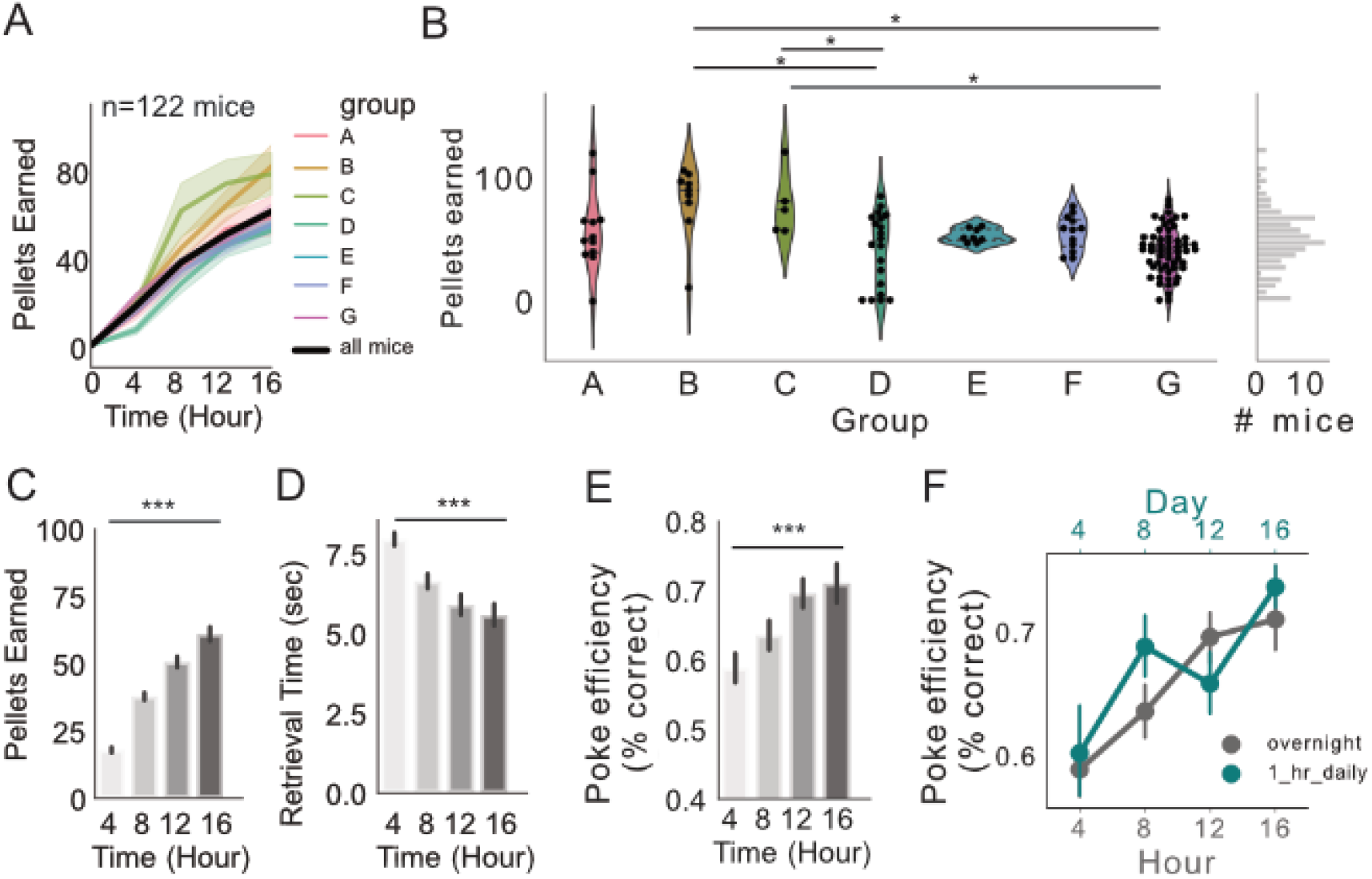
FR1 acquisition across seven research sites. **(A)** Pellets earned over first 16 hours of exposure to FED3 at seven research sites. **(B)** Scatterplots and kernel density estimation plots (KDE) showing pellets earned after 16 hours with FED3, effect of group (F_(6,115)_=6.4223, p<0.0001), significant post-hoc differences between groups B and D (p=0.001), B and G (p=0.001), C and D (p=0.030), and C and G (p=0.010). **(C)** Pellets earned across the session. Significant effect of time on average active poke count, p=0.0001, F(3, 393)=115.2. **(D)** Retrieval time across the session. Significant effect of time, p=0.0001, F(3,351)=14.02. **(E)** Poke efficiency across the session. Significant effect of time, p=0.0007, F(3,386)=5.79. Linear models with Tukey post-tests were conducted in panels **C-E**. **(F)** Poke efficiency across continuous 16 hour sessions (grey line, bottom x-axis, n=122 mice) and across a 16 days with 1 hr sessions each day (teal line, top x-axis, n=11 mice). 2-way ANOVA revealed significant effect of time (p<0.0001), no significant effect of group or interaction. F(7,500)=3.974.

A unique capability of FED3 is that is has a sensor for detecting the pellet itself. This enables time-stamping of when each pellet was removed for constructing feeding records, as well as measuring the time between the pellet dispense and removal, which we termed “retrieval time”. Retrieval time can be used as an index of learning, as it progressively decreased as animals gained experience with FED3 (Figure 5D). Accordingly, poke efficiency (% of pokes on the active port) increased over time, demonstrating that mice learned which poke resulted in a pellet (Figure 5E). To compare learning rates in overnight training to traditional daily operant sessions, we ran a new group of eleven mice for sixteen daily 1-hour FR1 sessions with FED3, returning them to the colony for the remainder of each day. Learning rates did not differ between mice exposed to sixteen daily 1-hour sessions, or one 16-hour over-night session (Figure 5F). This suggests that the acquisition of operant tasks depends on the cumulative time mice are exposed to FED3, and overnight training can speed up task acquisition.

### The effect of magazine training on acquisition of operant behavior

Based on prior literature (Steinhauer et al., 1976), we predicted that prior magazine training would also speed up learning about operant associations in an FR1 task. To test this, we exposed 24 mice to magazine training using the free feeding FED3 paradigm for 1-3 days prior to FR1 training (Figure 6A). This paradigm dispenses a pellet into the feeding well of FED3 and replaces it whenever it is removed, providing an automatic method for magazine training animals. This allowed mice to learn the location of the food source and associate food with the sound of the pellet dispenser operating and the pellet being dispensed, before commencing with FR1 training. We compared the performance of the 24 magazine trained mice to 122 mice from Figure 5 (which had not been magazine trained), and found that magazine training resulted in significantly higher (~2x) levels of pellet acquisition in the first night with the FR1 task (Figure 6B, C). Therefore, magazine training is recommended to speed up acquisition of nose-poking tasks.

**Figure 6.**
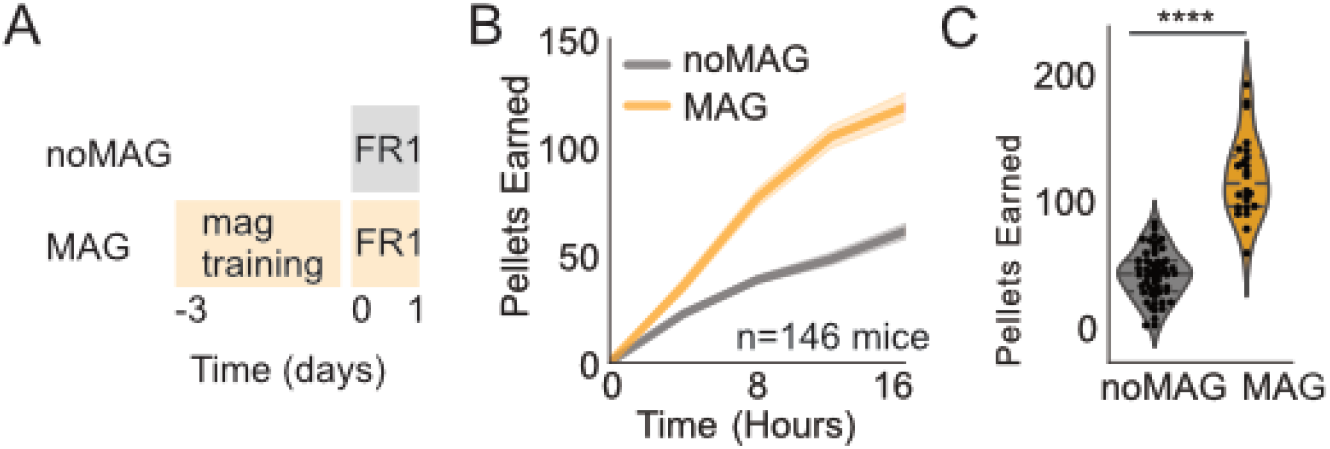
Effect of magazine training. **(A)** Schematic showing paradigm for magazine trained group (MAG) vs no magazine training (noMAG). **(B)** Active pokes over time during first exposure to FED3 FR1 between noMAG and MAG groups. **(C)** Scatterplots and KDE plots showing distribution of active poke counts at 16 hours (p=0.0001). Student’s t-test. N=146 mice.

### Optogenetic self-stimulation with FED3

FED3 was also designed to synchronize with other experimental equipment through its programmable output port. To demonstrate this capability, we programmed FED3 to control an LED for an intra-cranial self-stimulation task. The dorsal striatum is composed of two populations of output neurons, differentiated by their expression of the dopamine D1 or D2 receptors (Gerfen et al., 1990). Dorsal striatal neurons that express the D1 receptor are highly reinforcing when optogenetically stimulated (Kravitz et al., 2012). To demonstrate how FED3 can be used to perform optogenetic self-stimulation experiment, we used Cre-dependent viral expression to target channelrhodopsin-2 (ChR2) to D1R-expressing neurons in the dorsal striatum (Figure 7A). FED3 was programmed to trigger a 1 second train of 20Hz pulses of 475nm light upon each active nose-poke, and the session continued until they received 75 stimulations (Figure 7B). Three mice were run on this task, resulting in significantly greater active vs inactive pokes (Figure 7C, D), and demonstrating that FED3 can be used for optogenetic self-stimulation studies. The programmable output is attached to an on-board digital-to-analog converter chip, meaning that it can be programmed to output pulses of any voltage from 0-3V, allowing FED3 to represent multiple behavioral events with one output line.

**Figure 7.**
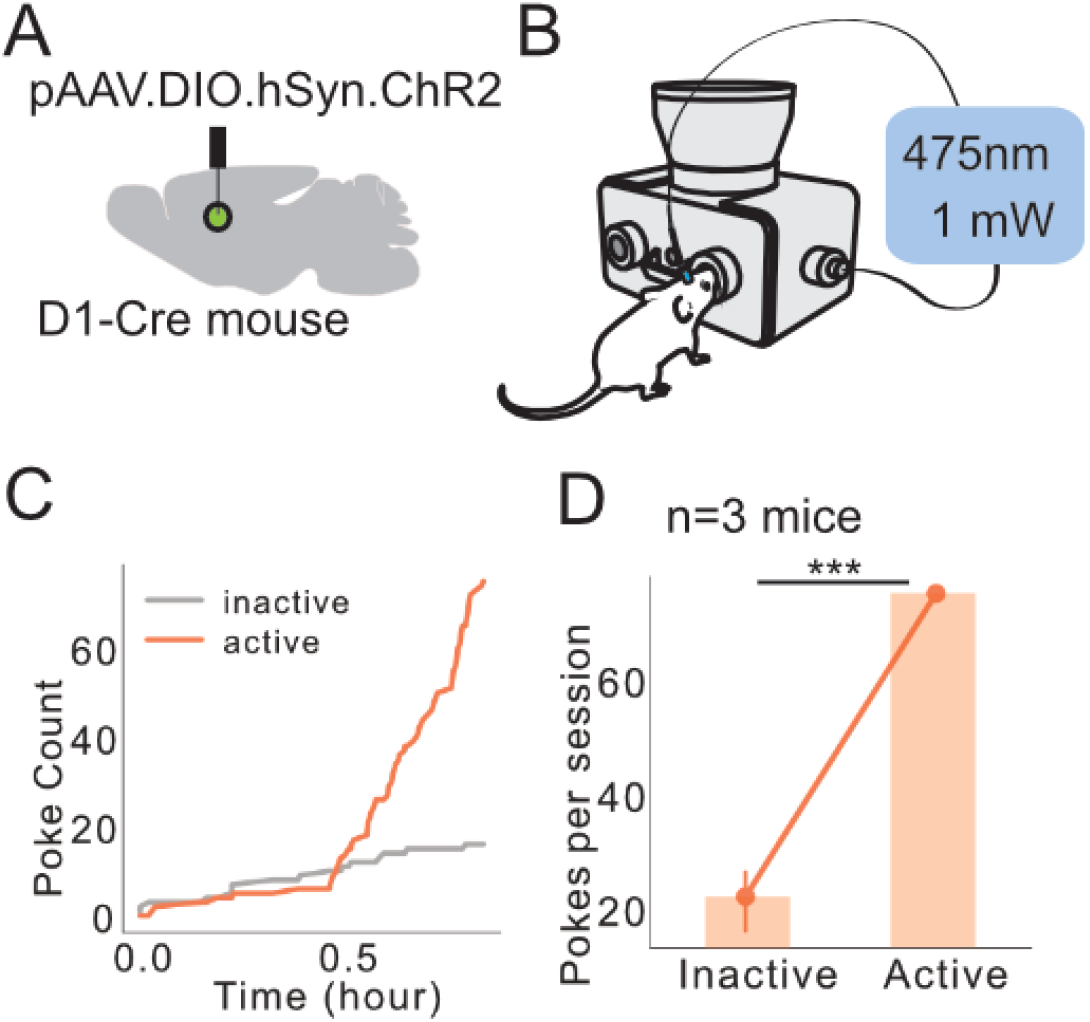
Self-stimulation of dMSNs using FED3. **(A)** Schematic showing Cre-dependent ChR2 injected into dorsal medial striatum with fiber optic implanted. **(B)** Schematic showing FED3 operant self-stim setup. **(C)** Example data from single mouse showing inactive and active pokes. (D) Mice poked significantly more on the active port (p=0.0002, Student’s t-test, n=3 mice).

## Discussion

Quantification of food intake is necessary to understand animal models of feeding disorders, as well as other medical conditions. However, food intake measurements are often completed using manual methods that are time consuming, error prone, and do not measure motivation. To automatically quantify food intake in rodent home cages, we previously published the Feeding Experimentation Device (FED), an open-source, stand-alone feeding device that can fit in a rack-mounted home-cage (Nguyen et al., 2016). FED records a time-series of pellets removed, which can be used to reconstruct feeding records. FED has been used by multiple research groups to understand feeding (Brierley et al., 2020; Burnett et al., 2019; Chen et al., 2020; Li et al., 2019; London et al., 2018). Since its original publication, we have redesigned the FED twice and here present the 3^rd^ version of FED, which revamps our original design and adds the ability to measure operant behavior, as well as a unique “angled” pellet dispenser mechanism that is resistant to jamming. We distributed the design for FED3 online and it has already been used to study how specific neural circuit manipulations alter food motivation (Mazzone et al., 2020; Sciolino et al., 2019; Vachez et al., 2020), how weight-loss alters food motivation (Matikainen-Ankney et al., 2020), and how food motivation is altered in a stress-susceptible mouse population (Rodriguez et al., 2020).

FED3 is a stand-alone solution for home-cage operant training, enabling researchers to understand not just total food intake, but also motivation for and learning about food rewards. FED3 has several unique benefits including: 1) FED3 is open-source and low cost. The FED3 electronics cost ~$150 and the housing is 3D printed. This is >10x cheaper than most commercial solutions for measuring food intake or testing operant behavior; 2) FED3 is self-contained and fits within traditional vivarium caging, allowing for measures of true “home cage” feeding without modifying the cage. Due to its small size it can also be placed inside of other equipment where wires might be impractical, such as within an indirect calorimetry system; 3) FED3 has a programmable output that allows it to easily synchronize with other equipment. This has already been used by multiple labs to synchronize the output of FED3 with fiber-photometry recordings (London et al., 2018; Mazzone et al., 2020), electrophysiological recordings (London et al., 2018), optogenetic stimulation (Vachez et al., 2020), and video tracking (Krynitsky et al., 2020; Li et al., 2019); and 4) FED3 is open-source with all design files and code freely available online. This enables users to modify the code and hardware to achieve new functionality.

The purpose of this manuscript is to demonstrate the utility of FED3 for feeding research. To this end, we demonstrate experiments that measured total food intake, operant responding, and optogenetic stimulation. We further highlight how the high temporal resolution enables meal pattern analysis across multiple days. Finally, we coordinated with 6 other research groups to compile a dataset of 122 mice across 7 research sites, all running the same experimental fixed-ratio 1 (FR1) program. We observed similar patterns of acquisition across all sites, demonstrating that FED3 can be used for multi-site research studies on feeding. Due to its low cost and open-source nature, we believe that multi-site studies with FED3 will be more feasible than with commercial equipment.

While FED3 has many strengths, it also has limitations. One limitation is that FED3 uses internal microSD cards to store data. While microSD cards are convenient, they are not ideal for large numbers of devices, where removing multiple cards can be cumbersome. Wireless data logging is a potential solution to this problem, although there are challenges to implementing this in rodent cages. Another limitation is that animals can “hoard” pellets from FED3, as animals can remove food without consuming it. In our experience, this seemed to be a trait that specific mice engaged in, but was rare (<10% of mice hoard pellets). Unfortunately, FED3 has no way to determine if a mouse consumes every pellet it eats so we recommend checking for pellet hoarding and accounting for this in experimental conclusions. A second limitation is that granular bedding can be kicked into the pellet well and interfere with pellet detection. To avoid this, we recommend using “iso-pad” bedding, or bedding pellets that are large enough to avoid this possibility. One final limitation of FED3 is that it currently has no way of identifying individual mice in group housed environments, so feeding records must be collected in singly housed mice. However, FED3 is open-source so it can be modified and improved with innovations from our group and the feeding community, and future iterations of the device may include methods for identifying multiple mice using radio-frequency identification (RFID) tags. Being able to run studies on group housed mice would greatly increase throughput and allow for the study of interactions between social behavior and feeding. We published the FED3 design as open-source and look forward to community contributions and modifications to enable new functionality and overcome these limitations.

## Materials and Methods

Key Resources Table.

Key Resources Table
| Reagent | Source | Identifier |
| --- | --- | --- |
| Software and Algorithms |  |  |
| Python 3.7 (Anaconda Distribution) | https://www.anaconda.com/ |  |
| Spyder 4.1 | https://github.com/spyder-ide/spyder/releases |  |
| Arduino IDE | https://www.arduino.cc/ |  |
| FED3_VIZ | https://github.com/earnestt1234/FED3_Viz |  |
| TinkerCAD | https://www.tinkercad.com/ |  |
| Other |  |  |
| FED3 – commercially assembled | https://open-ephys.org/fed3/fed3 | N/A |
| FED3 – open source | https://github.com/KravitzLabDevices/FED3 | N/A |
| 20mg Food pellets | https://www.testdiet.com/Diet-Enrichment-Products/Lab-Treat-Tablets-and-Pellets/index.html | N/A |

### Data and code availability

The FED3 device is open-source and design files and code are freely available online at: https://github.com/KravitzLabDevices/FED3. In addition, we have made all data and analysis code for this paper available at https://osf.io/hwxgv/. Any other request for data or code can be made to the corresponding author.

### Subjects

159 C57Bl6 mice were housed in a 12-hour light/dark cycle with ad libitum access to food and water except where described. Mice were provided laboratory chow diet (5001 Rodent Diet; Lab Supply, Fort Worth, Texas). All procedures were approved by the Animal Care and Use Committee at Washington University in St Louis, the National Institutes of Health, Williams College, Virginia Tech and Monash University.

#### Design and construction of FED3

Tutorial videos and other information on assembling FED3 are available at: https://github.com/KravitzLabDevices/FED3/wiki.

##### 3D design

The 3D parts for FED3 were designed with TinkerCAD (Autodesk). We have printed FED3 with a FDM printer in PLA (Sindoh 3DWox 1), as well as with a commercial SLS process in Nylon 12 (Shapeways). 3D files may need to be tweaked for specific printers, and editable design files are here: https://www.tinkercad.com/things/0QaiVw7KR3Y.

##### Electronics

Electronics design was completed using Autodesk Eagle version 9.3. FED3 is controlled by a commercial microcontroller (Adalogger Feather M0, Adafruit). This microcontroller contains an ATSAMD21G18 ARM M0 processor that runs at 48MHz, 256KB of FLASH memory, 32KB of RAM memory, an on-board microSD card slot for writing data, and up to 20 digital inputs/output pins for controlling other hardware. The microcontroller also contains a battery charging circuit for charging the internal 4400mAhr LiPo battery in FED3, which provides ~1 week of run-time between charges. Exact battery life depends on the behavioral program and how often the mouse interacts with FED3. Additional hardware on FED3 includes a motor for controlling the pellet delivery hopper, 8 multi-color LEDs for delivering visual stimuli, a small speaker for delivering audio stimuli, a screen for user feedback, and three infra-red beam-break sensors. Two of these sensors are used as “nose-poke” sensors to determine when the mouse pokes, while the third is used to detect the presence and removal of each food pellet. Finally, FED3 contains a programmable output connected to a 10-bit digital-audio converter circuit, enabling full analog output of arbitrary voltages from 0-3.3V. This can be used to synchronize FED3 with external equipment via digital pulses or analog signals. There are two variants of the electronics for FED3: An older design is referred to as the “DIY FED3” which can be assembled by hand using commercially available components (https://hackaday.io/project/106885-feeding-experimentation-device-3-fed3), whereas a newer design contains small improvements to the electronics but requires professional electronics assembly. Both designs run the same code and have identical functionality.

##### Firmware

The code for FED3 is written in the Arduino language and is fully open-source. The standard code that we provide with FED3 contains 12 programs, covering multiple common behavioral paradigms, including free feeding, time-restricted free-feeding, fixed-ratio (FR1), and progressive ratio (PR) operant tasks, as well as an optogenetic self-stimulation mode that delivers trains of pulses for controlling a stimulation system. We also provide an Arduino library that simplifies control of FED3 hardware to simplify writing of custom programs.

### Behavioral testing with FED3

We demonstrate multiple operational modes for FED3 to highlight a range of functionality and applications. The following programs are included with the standard FED3 code:

#### Free-feeding

In the free-feeding mode (Figure 2), a 20mg pellet is dispensed into the feeding well and monitored with a beam-break. When the pellet is removed, the time-stamp of removal, and the latency to retrieve the pellet are logged to the internal microSD card and a new pellet is dispensed. For the free-feeding data in this paper, 10 mice were singly housed and the FED3 device was placed in their cage for 6 days on a 12/12 on/off light cycle. FED3 devices were checked for functionality each day but mice were otherwise undisturbed.

#### Fixed-ratio 1 (FR1)

In FR1 mode, FED3 logs the time-stamps of each “nose-poke” event to the internal storage. When the mouse activates the left nose-poke the FED3 delivers a combined auditory tone (4kHz for 0.3s) and visual (all 8 LEDs light in blue) stimulus, and dispenses a pellet. While the pellet remains in the well both pokes remain inactive to prohibit multiple pellets piling up in the well. When the pellet is removed, the time-stamp of removal, and the latency to retrieve the pellet are logged. For the data in this paper, the same 10 mice that completed the Free-feeding experiment were transitioned to an FR1 program, and FR1 data was recorded for an additional 6 days.

##### Optogenetic stimulation

Viral infections of male and female Drd1-Cre mice were conducted under 0.5-2.5% anesthesia on a stereotaxic apparatus. Using a Nanoject injector, 500 nL of AAV2-DIO-hSyn-ChR2 was injected bilaterally (500 nL/ hemisphere) in the dorsomedial striatum. Optical fibers were then implanted in the same region. After letting ChR2 express for four weeks, animals were pre-trained on an FR1 schedule for pellets overnight in their home cage. Two days later mice were placed in a box with a FED3 device connected to an LED driver. The active poke for this paradigm was on the opposite side to the active poke of the pre-training session. During the self-stimulation session, when the mouse poked on the active nose-poke, it received a 1 second train of 1 mW 475 nm light at 20 Hz. Sessions were run for 60 minutes.

### Data Analysis

CSV files generated by FED3 were processed and plotted with custom python scripts (Python, version 3.6.7, Python Software Foundation, Wilmington, Delaware). All data and scripts are available on Open Science Framework (https://osf.io/hwxgv/). Visualization was also completed using FED3VIZ GUI to generate plots. FED3VIZ was written in Python’s standard library for developing GUIs (tkinter). FED3VIZ is a custom open-source graphical program for analyzing FED3 data. FED3VIZ code, version history, installation instructions, and user manual are available on GitHub (https://github.com/earnestt1234/FED3_Viz). FED3VIZ offers plotting and data output for visualizing different aspects of FED3 data, including pellet retrieval, poke accuracy, pellet retrieval time, delay between consecutive pellet earnings (interpellet intervals), meal size, and progressive ratio breakpoint. Based on inter-pellet interval histograms (Figure 3), we defined meals as pellets eaten within 1-minute of each other. In addition, we defined a minimum size of 0.1g (5 pellets) to be counted as a meal.

#### Statistics

Bartlett’s test for equal variances was performed; one- or two-way ANOVAs with Tukey post-tests were used to compare groups with equal variance (P>0.05) where appropriate. Linear mixed effects models were used to analyze groups with significantly difference variances (p<0.05). P-values <0.05 was considered significant. Statistical tests to compare means were run using statsmodels module in python (Seabold and Perktold, 2010). Data sets are presented as mean +/− SEM. Numbers of animals per experiment is listed as n=number of animals. Linear regression was used to determine correlative relationships. T-tests or Mann Whitney U tests were used to compare the means of two groups for parametric or nonparametric data distributions, respectively.

## Author contributions

**Bridget A. Matikainen-Ankney**: Methodology, Data Curation, Formal analysis, Writing **Thomas Earnest**: Methodology, Software **Mohamed Ali**: Methodology, Software **Eric Casey**: Validation, Investigation **Amy K. Sutton**: Investigation, **Alex Legaria**: Investigation, **Kia Barclay**: Review & Editing, **Laura B. Murdaugh**: Investigation, **Makenzie R. Norris**: Investigation, **Yu-Hsuan Chang**: Investigation, **Katrina P. Nguyen**: Methodology, **Eric Lin**: Software, **Alex Reichenbach**: Investigation, **Rachel E. Clarke**: Investigation, **Romana Stark**: Investigation, **Sineadh M. Conway**: Investigation, **Filipe Carvalho**: Methodology, **Ream Al-Hasani**: Resources, Review & Editing, Supervision **Jordan G. McCall**: Resources, Review & Editing, Supervision, **Meaghan C. Creed**: Resources, Review & Editing, Supervision, **Victor Cazares**: Resources, Review & Editing, Supervision, **Matthew W. Buczynski**: Resources, Review & Editing, Supervision, **Michael J. Krashes**: Resources, Review & Editing, Supervision, **Zane Andrews**: Resources, Review & Editing, Supervision, and **Alexxai V. Kravitz**: Methodology, Software. Resources, Writing, Review & Editing, Supervision, Funding acquisition

## Acknowledgements

Funding for developing FED was provided from the National Institutes of Health Intramural Research Program at NIDDK (AVK) and the Washington University Diabetes Research Center Pilot and Feasibility Grant (AVK).

